# Regulation of brain development by the Minibrain/Rala signaling network

**DOI:** 10.1101/2024.05.10.593605

**Authors:** Melissa Brown, Erika Sciascia, Ken Ning, Wesam Adam, Alexey Veraksa

**Affiliations:** Department of Biology, University of Massachusetts Boston, Boston, MA 02125, USA

## Abstract

The human dual specificity tyrosine phosphorylation regulated kinase 1A (DYRK1A) is implicated in the pathology of Down syndrome, microcephaly, and cancer, however the exact mechanism through which it functions is unknown. Here, we have studied the role of the *Drosophila* ortholog of DYRK1A, Minibrain (Mnb), in brain development. The neuroblasts (neural stem cells) that eventually give rise to differentiated neurons in the adult brain are formed from a specialized tissue in the larval optic lobe called the neuroepithelium, in a tightly regulated process. Molecular marker analysis of *mnb* mutants revealed alterations in the neuroepithelium and neuroblast regions of developing larval brains. Using affinity purification-mass spectrometry (AP-MS), we identified the novel Mnb binding partners Ral interacting protein (Rlip) and RALBP1 associated Eps domain containing (Reps). Rlip and Reps physically and genetically interact with Mnb, and the three proteins may form a ternary complex. Mnb phosphorylates Reps, and human DYRK1A binds to the Reps orthologs REPS1 and REPS2. Furthermore, Mnb engages the small GTPase Ras-like protein A (Rala) to regulate brain and wing development. This work uncovers a previously unrecognized early role of Mnb in the neuroepithelium and defines the functions of the Mnb/Reps/Rlip/Rala signaling network in brain development.

**Significance statement:** The kinase Minibrain(Mnb)/DYRK1A regulates the development of the brain and other tissues across many organisms. Here we show the critical importance of Mnb within the developing neuroepithelium. Advancing our understanding of Mnb function, we identified novel protein interactors of Mnb, Reps and Rlip, which function together with Mnb to regulate growth in *Drosophila melanogaster*. We also identify and characterize a role for the small GTPase Rala in Mnb-regulated growth and nervous system development. This work reveals an early role of Mnb in brain development and identifies a new Mnb/Reps/Rlip/Rala signaling axis.

## Introduction

The kinase Minibrain (Mnb) and its orthologs control the development of the brain and other tissues in *Drosophila* and mammals (1–3). In *Drosophila,* both loss and gain of *mnb* function can result in abnormal development. Reduction of *mnb* function results in decreased brain and wing size, disrupted wing vein patterning, abnormal visual and food intake behavior, and defects in dendrite development (1, 3–6). Gain of Mnb is associated with a shortened lifespan, impaired neuronal function, and accelerated decline in motor performance (7). Likewise, the homolog of Mnb, dual specificity tyrosine phosphorylation regulated kinase 1A (DYRK1A), is critical for mammalian development and exhibits both loss and gain of function phenotypes (8–12). In humans, the *DYRK1A* gene is located on chromosome 21, and through studies in mouse models, *DYRK1A* copy number has been shown to play a causative role in Down syndrome neurodevelopmental defects (11, 13). Loss of function mutations in *DYRK1A* that lead to decreased DYRK1A protein levels result in primary microcephaly and developmental delays, reminiscent of *mnb* mutant phenotypes (8, 12, 14). Given its disease phenotypes, it is clear that DYRK1A plays a critical role in neuronal development and physiology. DYRK1A has also been implicated in the control of the cell cycle and cell proliferation. Depending on the cellular context, DYRK1A can either promote or inhibit cell proliferation, and has both oncogenic and tumor suppressor characteristics (15, 16).

*Drosophila* is a useful model system to study brain development, given the availability of genetic tools and similarities in neurogenesis between flies and mammals. In both *Drosophila* and mammalian brain development, there are stem cell populations that divide symmetrically to amplify a progenitor population, which then transitions to asymmetric divisions (17–19). In the optic lobe of the fly third instar larval brain, the symmetrically dividing population is the neuroepithelium (NE), which gives rise to asymmetrically dividing stem cells, known as the neuroblasts (NBs) (20). Reduction of *mnb* function disrupts NB divisions and induces cell cycle defects, which ultimately leads to apoptosis and a decrease in the adult brain size (21, 22). However, the molecular mechanisms of how Mnb controls the formation and functions of neural stem cells are incompletely understood.

Here we uncovered a previously unknown early developmental role of Mnb in controlling the transition between the neuroepithelium and neuroblasts. Using affinity purification-mass spectrometry (AP-MS), we identified the protein interaction network of Mnb and characterized RALBP1 Eps domain containing (Reps) and Ral interacting protein (Rlip) as novel binding partners of Mnb. We show that Reps and Rlip function together with Mnb to regulate the growth and development of the brain and the wing. We also determined that Mnb phosphorylates Reps, and that the physical interaction between Mnb and Reps is conserved across their human orthologs, DYRK1A and REPS1/2. Lastly, we investigated a connection between Mnb and the small GTPase Ras-like protein A (Rala), whose function is closely related to that of Rlip, and found that Mnb and Rala jointly control brain and wing growth. Collectively, this work establishes an early function of Mnb in controlling neuroepithelium functions and identifies the Mnb/Reps/Rlip/Rala signaling axis that regulates brain development.

## Results

### Mnb regulates neuroepithelium development

Previous work investigating the role of Mnb in larval brain development primarily focused on neuroblasts and their progeny in the central brain (CB) (21, 22). Given the significant size reduction of the adult optic lobes observed in *mnb* mutants (3, 4), we investigated Mnb’s role during earlier stages of optic lobe development. The optic lobes in the adult brain are derived from NBs that are tightly packed along the surface of the larval brain in a region called the outer proliferation center (OPC) (Fig. 1A) (23, 24). Unlike NBs located in the CB, OPC NBs arise during larval development from neuroepithelial (NE) cells which are located directly adjacent to NBs. The NE cells must undergo a highly regulated transition from NE to NB. This transition is mediated by a proneural wave that is controlled by various signaling pathways including EGFR and Notch (Fig. 1A) (25).

**Figure 1.**
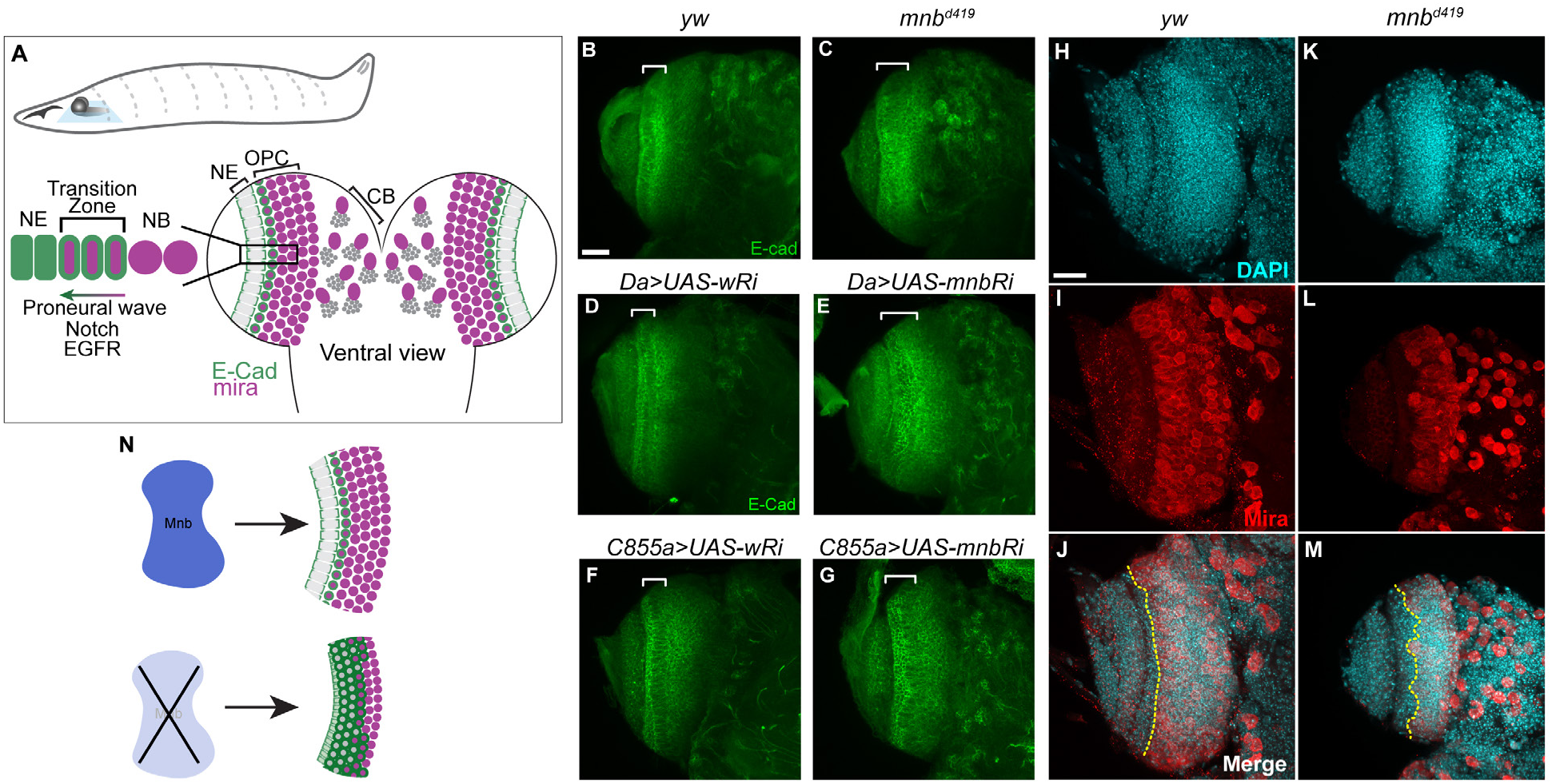
Mnb is required for neuroepithelium and neuroblast organization. (A) Top: overall view of third instar larva with central nervous system shown. A blue square indicates imaging plane (ventral view). Bottom: Neuroepithelium (NE) to neuroblast (NB) transition is mediated by Notch and EGFR signaling. Adapted from (30, 53). (B-G) Confocal maximum intensity projections of larval brains of the indicated genotypes immunostained for E-cad. Brackets indicate the E-cad expression domain corresponding to the NE. (H-M) Confocal maximum intensity projections of larval brains of the indicated genotypes immunostained for Miranda (Mira) shown in red, with DAPI signal shown in cyan. Dotted line marks a boundary between the NE and the OPC. Scale bars, 25 µm. (N) Schematic representing NE/NB outcomes in WT and *mnb* mutant conditions.

We used the mutant allele *mnb^d419^* (5) which deletes part of the kinase domain in the Mnb protein (Fig. S1D) and results in smaller 3^rd^ instar larval brains in hemizygous male animals (3). These hemizygous larvae formed pharate adults that died during eclosion, which is a more severe phenotype than the one observed for the kinase-inactive *mnb^1^* mutant allele that is homozygous viable (4) (6). Heterozygous female flies containing one copy of *mnb^d419^* and one copy of *mnb^1^* were viable (Fig. S1). To ensure that *mnb^d419^* male lethality was not due to a mutation outside the *mnb* locus, we crossed *mnb^d419^* to two lines carrying duplications of the 1^st^ chromosome containing the *mnb* gene and found that these lines rescued *mnb^d419^* lethality (Fig. S2). These results demonstrate that the *mnb^d419^* allele is a strong loss of function mutation in *mnb*.

To visualize the NE, we stained for the epithelium marker E-cadherin (E-cad) (26, 27). In wild-type animals, E-cad staining revealed a distinct NE organization with columnar epithelial cells expressing E-cad at their basal and apical sides (Fig. 1B, D, F, bracketed region). A few dividing epithelium cells can be identified as the rounded cells within the NE band and only a few cells within the OPC are E-cad positive. In contrast, the NE organization in *mnb^d419^* was disrupted, resulting in less columnar cells and expansion of E-cad staining into the NB region of the OPC (Fig. 1C). Larval brains expressing *UAS-mnb RNAi* (*Ri*) under the control of the ubiquitous *da-GAL4* driver or the NE-specific driver *c855a-GAL4* showed a similar phenotype, compared to larval brains expressing *UAS-wRi* as a control (Fig. 1D-G). These results suggest that Mnb is required for the proper organization and development of the NE in the optic lobe.

Given that NE cells give rise to NBs, we assessed the organization of NBs through staining for the NB marker Miranda (Mira) (28, 29). Wild-type larval brains showed a distinct band of tightly packed Mira positive NBs that make up the OPC (Fig. 1I). The boundary between Mira positive NBs and the Mira negative NE cells was straight and smooth in wild-type brains (Fig. 1H-J, dotted line). In contrast, the boundary between NE and NB in *mnb^d419^* mutant brains was jagged and uneven (Fig. 1K-M), suggesting there may be Mira negative, E-cad positive cells persisting into the OPC. This disruption suggests a defect in NE to NB transition, a process that is tightly regulated by Notch and EGFR signaling (30).

Increased apoptosis was observed in *mnb^d419^* larval brains, based on cleaved Death caspase-1 (Dcp-1) staining (Fig. S3), in agreement with previously reported results using other *mnb* mutants (21). Based on the significant increase in Dcp-1 intensity within the OPC and the mis-localization of the E-cad and Mira markers within this same region, we speculate that these cells are mis-specified and undergo programmed cell death, ultimately resulting in a smaller adult brain. Collectively, these data have uncovered a previously unknown early role of Mnb in regulating NE function (Fig. 1N).

### Reps and Rlip interact with Mnb

To further characterize the mechanisms through which Mnb regulates brain development, we identified Mnb interacting proteins using AP-MS in embryos from endogenously tagged Mnb-tRFP (3) and control flies. Peptide counts were analyzed by Significance Analysis of INTeractome (SAINT), and proteins with interaction probability score above 0.8 were considered significant (31). The resulting interaction network of significant hits was generated in Cytoscape using the STRING protein interaction database and clustered using the clusterMaker MCL Cluster app (Fig. S4, see Materials and Methods) (32). Apart from Mnb itself, one of the top hits was the known Mnb interactor Wings apart (Wap) (1, 3) (Fig. 2A). We also found Regulator of eph (Reph) which was previously identified as a Mnb interactor in the *Drosophila* Protein Interaction Map (33) (Fig. 2A). Identification of these known interactors validates our AP-MS approach. Gene Ontology (GO) analysis of clusters showed several significantly enriched categories such as nuclear import (FDR=4.9E-13), ribosome biogenesis (FDR=8.01E-21), and endocytosis (FDR=1.64E-8) (Fig. S4).

**Figure 2.**
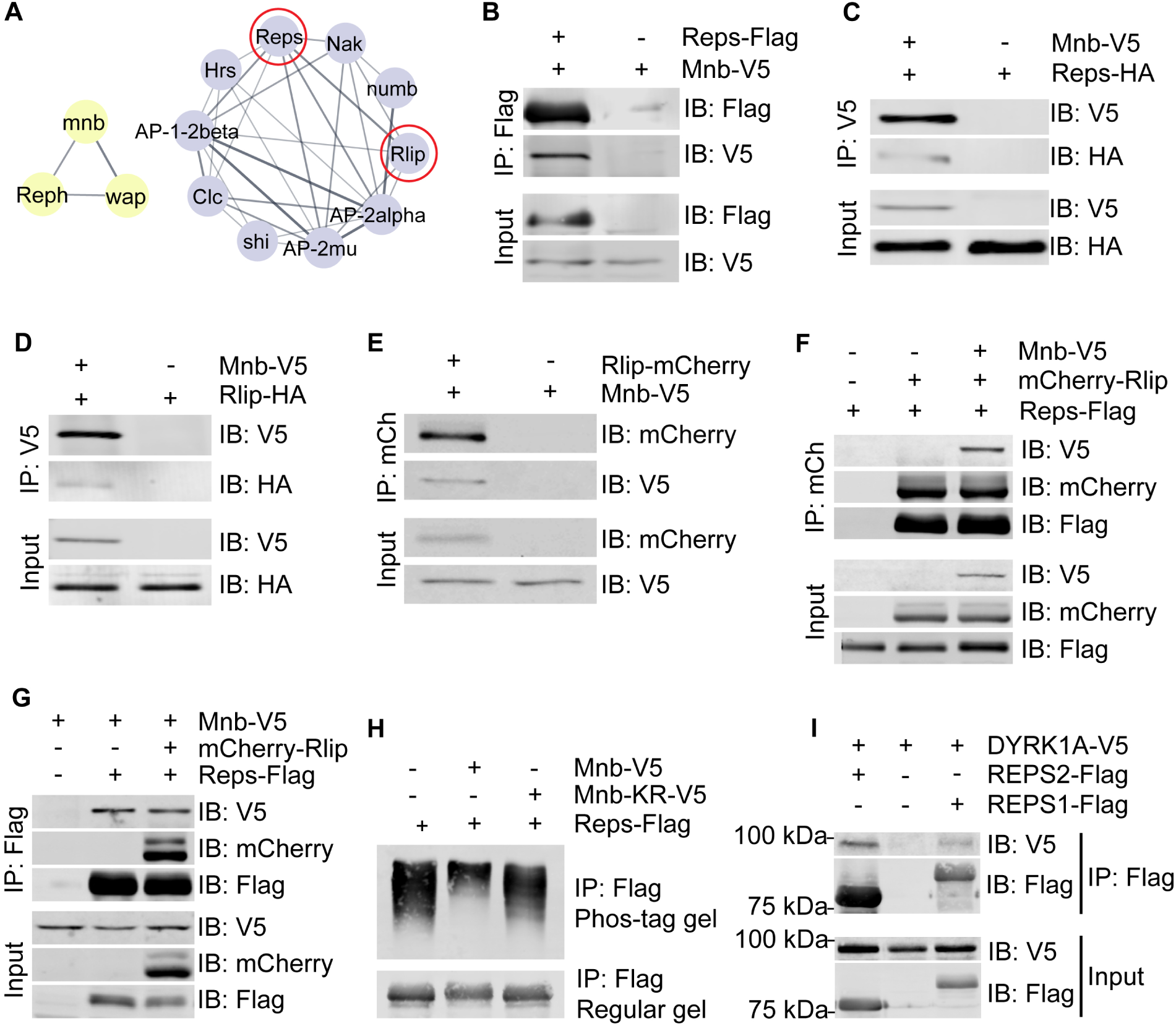
Mnb interacts with Reps and Rlip. (A) Network clusters identified in Mnb AP-MS include known Mnb interactors Wap and Reph (yellow) and a highly interconnected cluster of endocytosis related proteins (purple). (B-G) Co-immunoprecipitation (co-IP) experiments in *Drosophila* S2 cells showing the binding interactions among Mnb, Reps, and Rlip using the indicated protein combinations and IP directions. Mnb showed pairwise interactions with Reps (B, C) and Rlip (D, E), and a ternary interaction with Reps and Rlip (F, G). (H) Phos-tag analysis of Reps phosphorylation by Mnb. Bottom: regular SDS/PAGE gel. Mnb-KR, a kinase dead version of Mnb (Mnb^K193R^ mutant)(1). (I) Co-IP of DYRK1A and REPS1/2 in HEK293T cells through IP of REPS1/2.

The highly enriched complex of endocytic regulators (Fig. 2A) was interesting due to its potential impact on signaling, and we pursued characterization of two novel Mnb interactors from that cluster, RALBP1 associated Eps domain containing (Reps) and Ral interacting protein (Rlip). In mammals the orthologs of these proteins, REPS1/2 and ralA binding protein 1 (RALBP1), interact with activated small GTPase RalA and AP2 components to regulate receptor endocytosis (34). RALBP1 also has mitosis-specific roles and serves as an essential adaptor between Cdk1 and Epsin, enabling Cdk1-mediated phosphorylation of Epsin which renders it endocytosis-incompetent and functions as a molecular off switch for endocytosis during mitosis (35).

Co-immunoprecipitation (co-IP) of overexpressed tagged proteins confirmed the binding of Reps and Rlip to Mnb in *Drosophila* S2 cells (Fig. 2B-E). When expressed together, both Reps and Mnb can interact with Rlip, suggesting that these three proteins may form a ternary complex (Fig. 2F-G). Given the canonical role of Mnb as a kinase (3, 4, 36), we investigated whether Mnb could phosphorylate Reps and Rlip. Phos-tag gel analysis of Reps revealed a mobility shift upon co-expression with wild type Mnb but not kinase dead Mnb (Fig. 2H), suggesting that Mnb is sufficient to phosphorylate Reps and Mnb kinase activity is required for this phosphorylation. These findings validate Reps and Rlip as bona fide Mnb protein interactors.

In mammals, REPS1/2 and RALBP1 also physically interact (37, 38) but their interaction with the Mnb ortholog DYRK1A has not been reported. We found that REPS1 and REPS2 co-immunoprecipitated with DYRK1A in cultured HEK293T cells (Fig. 2I), whereas RALBP1 did not co-IP with DYRK1A (data not shown). These data show that the Mnb/Reps interaction is conserved, suggesting a possible involvement of DYRK1A in the regulation of REPS1/2 functions in human cells.

### Mnb promotes cytoplasmic Rlip localization

To probe the role of Mnb in the Mnb-Reps-Rlip complex, we investigated the localization patterns of these proteins in cultured *Drosophila* S2 cells. Mnb localized to the cytoplasm in a diffused pattern with a few distinct puncta in each cell (Fig. 3A-A’’). Rlip was found mostly in the nucleus, where it formed puncta, but was also occasionally distributed throughout the cell (Fig. 3B-B’’, quantified in Fig. 3F). Reps localized in a diffused cytoplasmic pattern (Fig. 3D-D’’). Co-expression of Mnb and Rlip resulted in subcellular redistribution of Rlip from mostly nuclear to primarily cytoplasmic or uniform localization (Figs. 3C-C’’’ and 3F). Mnb is therefore sufficient for retaining Rlip in the cytoplasm. This is significant because other Rlip interactors such as Rala are also localized in the cytoplasm, and Mnb may thus be promoting Rlip’s interaction with those factors. Reps localization remained unchanged upon co-expression with Mnb (Fig. 3E-E’’’).

**Figure 3.**
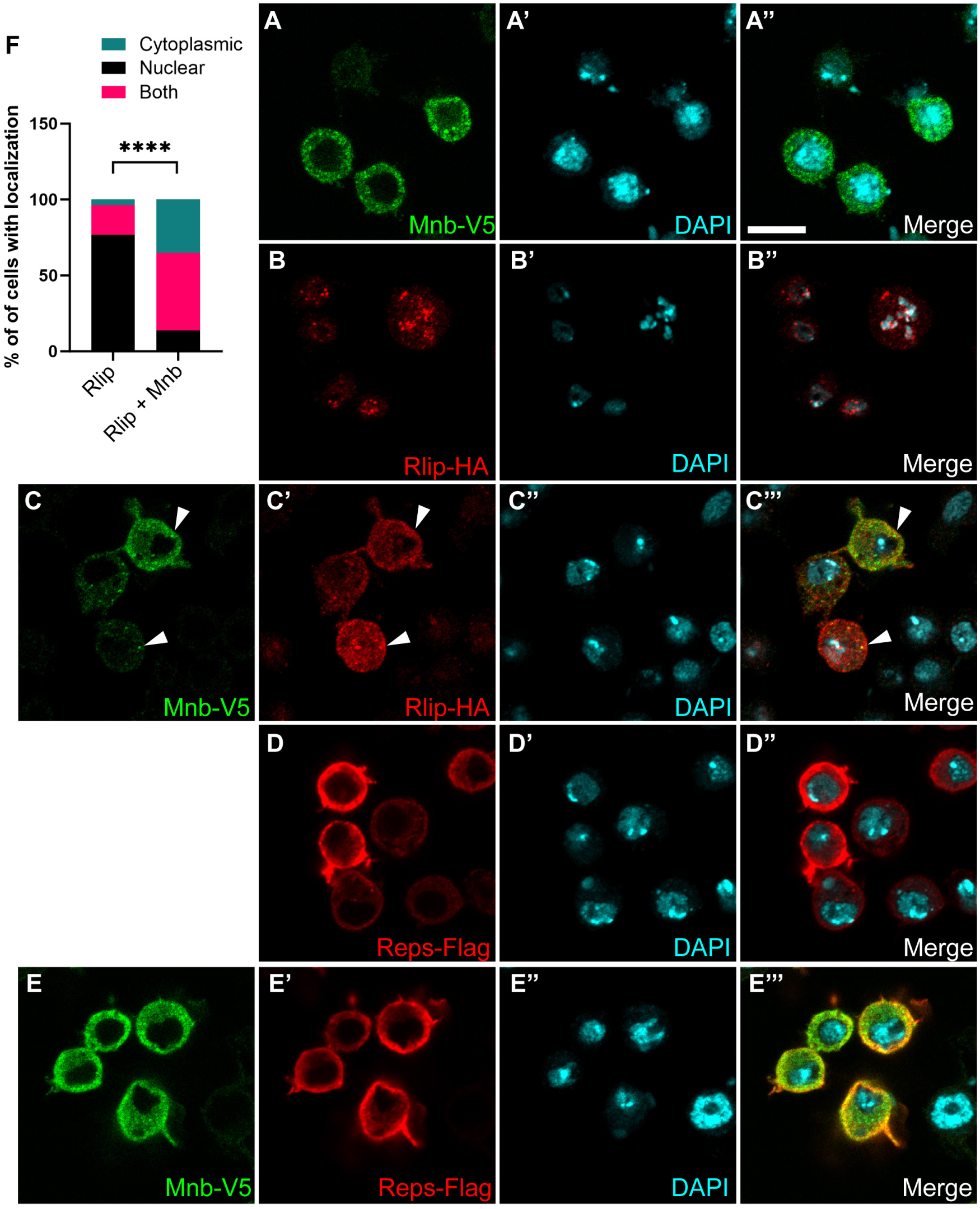
Mnb promotes cytoplasmic localization of Rlip. (A-E’’’) Confocal images of immunostained *Drosophila* S2 cells expressing Mnb-V5 (A-A”), Rlip-HA (B-B”), Mnb-V5 with Rlip-HA (C-C’’’), Reps-Flag (D-D”), and Mnb-V5 with Reps-Flag (E-E’’’). Arrowheads indicate Mnb-Rlip co-localization in cytoplasmic puncta (C-C’’’). Scale bar, 10 µm. (F) Quantification of Rlip localization in (C, D). Statistics calculated by chi-squared analysis, n>100 for each condition, **** *p* ≤ 0.0001.

### Reps and Rlip work together with Mnb to regulate wing growth

To determine whether Mnb, Reps, and Rlip are involved in common developmental processes, we tested the effects of knockdown of *Rlip* and *Reps* on the known *mnb* small wing phenotype. Knockdown of *mnb* with the wing-specific driver *MS1096-GAL4* resulted in a smaller wing than the control, consistent with previous observations (1, 3) (Fig. 4A, D, G). The knockdown of *Reps* did not alter wing growth but knockdown of *Rlip* resulted in a smaller wing than the control (Fig. 4A-C, G). Double knockdown of either *Reps* or *Rlip* in conjunction with *mnb* led to a further reduction in wing size, compared to the knockdown of *mnb* alone (Fig. 4D-G). These results suggest that Mnb, Reps, and Rlip work together to regulate wing growth.

**Figure 4.**
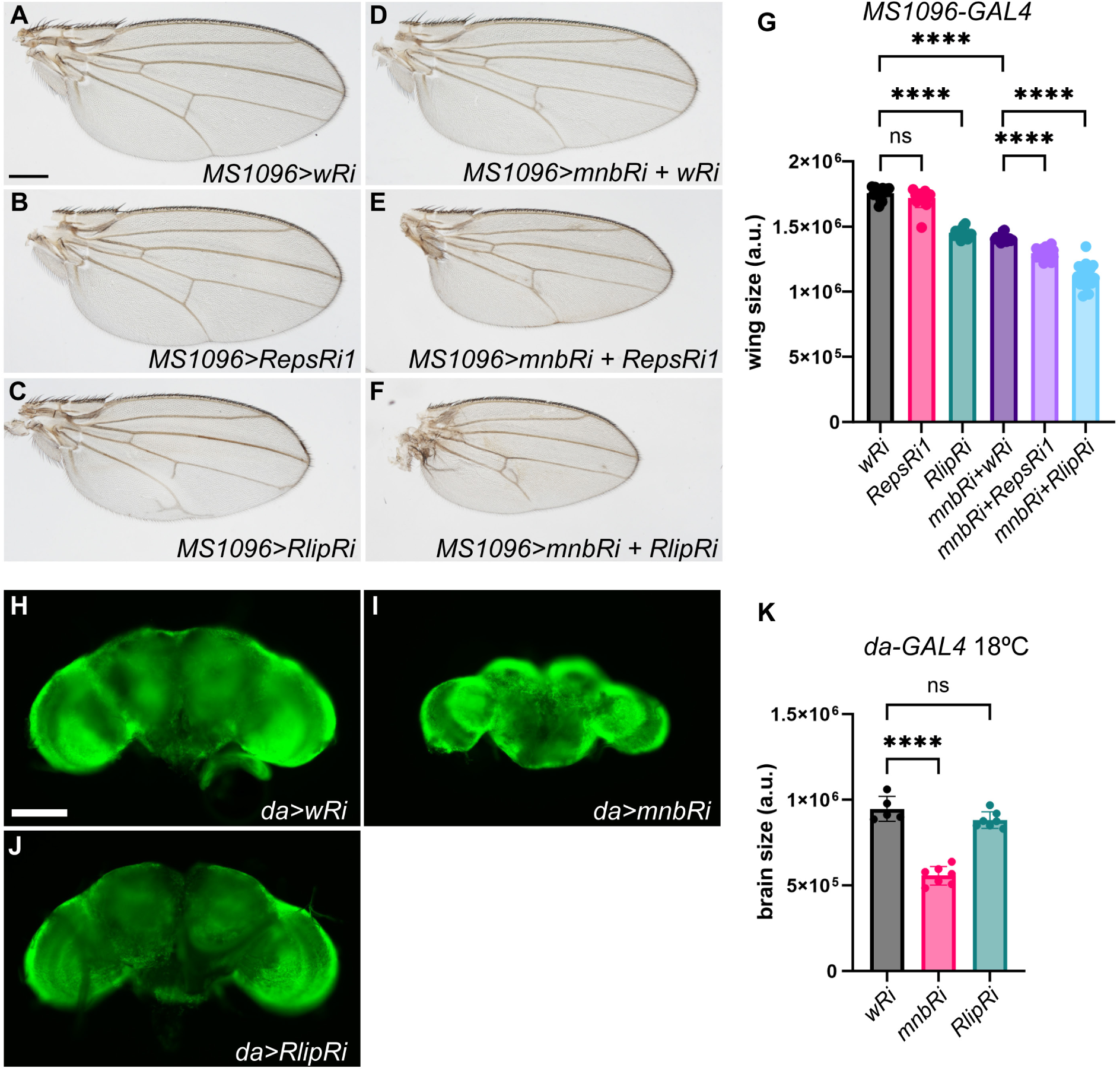
Mnb, Reps and Rlip work together to control wing and brain growth. (A-F) Wings from adult female flies expressing the indicated RNAi (Ri) transgenes using the wing-specific *MS1096-GAL4* driver. (G) Quantification of the wing areas shown in (A-F) (n≥14 for each genotype). \*\*\**p*<0.0001. *p* value calculated using ANOVA, error bars represent SD. Scale bar in (A), 300 µm. Adult brains from female flies grown at 18 °C (H-J) expressing the indicated RNAi transgenes. (K) Quantification of brain size shown in (H-J) (n ≥ 5 for each genotype). Brain size was measured as total area. DAPI signal is shown in green. \*\*\*\**p*<0.0001, *p* value was calculated using ANOVA. Error bars represent SD. Scale bar in (H), 100 µm.

### Rlip functions together with Mnb to regulate brain growth

Given the critical role of Mnb in brain development (3, 36), we asked whether Reps and Rlip also regulate brain growth. Knockdown of *mnb* with the ubiquitously expressed driver *da-GAL4* led to a decrease in adult brain size, particularly in the optic lobes (Fig. 4H-I). Knockdown of *Reps* with *da-GAL4* resulted in a larger brain than the control (Fig. S5A, C), however double knockdown of *Reps* and *mnb* resulted in a brain that was similar in size to that of *mnb* knockdown alone (Fig. S5A-E). This outcome suggests that Mnb was epistatic to Reps in this assay. Knockdown of either *Rlip* alone or *Rlip* in combination with *mnb* was lethal at 25 °C. However, when shifted to 18 °C, *RlipRi* flies were viable and did not show an altered brain size, compared to control flies also grown at 18 °C (Fig. 4J, K). *da-GAL4* driven knockdown of *mnb* at 18 °C resulted in smaller brains than the control as expected (Fig. 4I, K), however double knockdown of *mnb* and *Rlip* was lethal even at 18 °C, suggesting a synergistic effect. The results of this experiment suggest a model whereby Mnb and Rlip function together to control *Drosophila* development, and the requirement for Rlip becomes apparent in the sensitized genetic background of reduced *mnb* function.

### Mnb engages Rala for growth control

Rlip is a functional partner of the small GTPase Ras-like protein A (Rala) in the regulation of endocytosis and exocytosis of signaling receptors (38, 39). The Rlip ortholog preferentially interacts with activated GTP-bound Rala in *S. cerevisiae* (39). We compared Rlip interactions with wild type Rala and a constitutively active variant Rala^G20V^ (40). In *Drosophila* S2 cells, Rala^G20V^ bound to Rlip better than wild type Rala (Fig. 5A), confirming that *Drosophila* Rlip also preferentially interacts with activated Rala.

**Figure 5.**
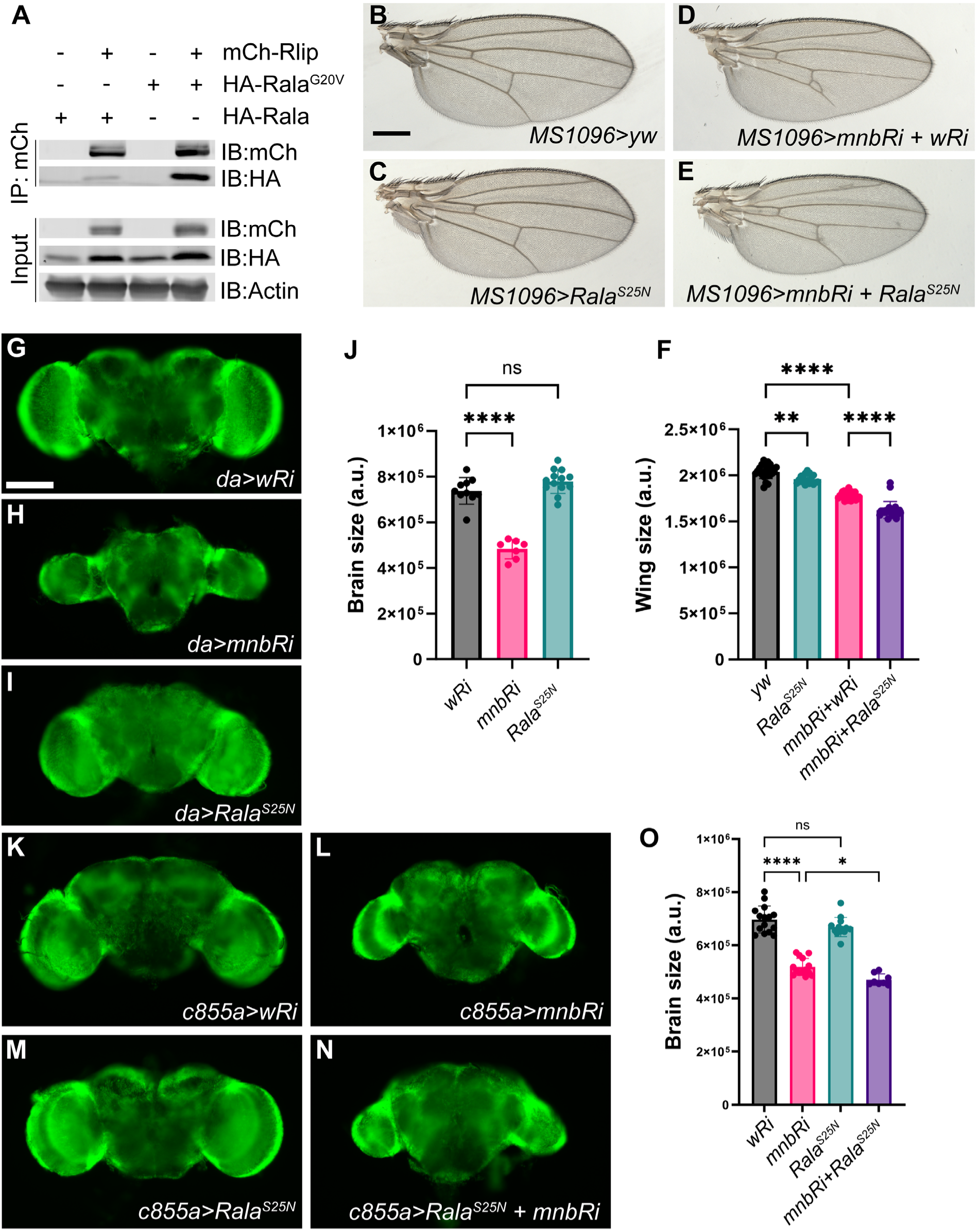
Mnb functions with Rala to regulate wing and brain development. (A) Co-IP of Rlip, wild type Rala, and Rala^G20V^ in *Drosophila* S2 cells through IP of Rlip. (B-E) Wings from adult female flies expressing the indicated transgenes using the *MS1096-GAL4* driver. A cross to *yw* (B) was used as a control. (F) Quantification of the wing areas shown in (B-E) (n≥19 for each genotype). Scale bar in (B), 300 µm. (G-I) Adult brains from female flies expressing the indicated RNAi (Ri) transgenes the ubiquitous *da-GAL4* driver. Scale bar in (G), 100 µm. (J) Quantification of the brain areas shown in (G-I) (n≥7 for each genotype). (K-N) Adult brains from female flies expressing the indicated transgenes using the NE-specific *c855a-GAL4* driver. (O) Quantification of brain size (area) shown in K-N (n≥8 for each genotype). \*\*\*\**p*<0.0001, \*\**p*<0.001, \**p*<0.01, *p* value calculated using ANOVA, error bars represent SD.

Given this connection between Rlip and Rala, we tested whether *Rala* and *mnb* interact genetically. We used a dominant-negative form of Rala (Rala^S25N^) that contains a serine to asparagine substitution at position 25 and as a result has a high affinity for GDP and cannot be activated (40). In the wing, overexpression of Rala^S25N^ slightly reduced wing size (Fig. 5B, C, F). Knockdown of *mnb* in conjunction with Rala^S25N^ overexpression reduced wing size more than either of these conditions alone (Fig. 5B-F). This suggests that Mnb and Rala cooperate to regulate wing growth. We also investigated the genetic interaction between Rala and Mnb in the adult brain. Interestingly, Rala^S25N^ overexpression phenocopied *Rlip-RNAi* in combination with *mnb-RNAi*: on its own, expression of Rala^S25N^ did not significantly alter adult brain size; however, when combined with *mnb* knockdown, this combination was lethal and adult brains could not be analyzed (Fig. 5G-J).

To determine whether Rala and Mnb function together in the NE, we used the NE-specific driver *c855a-GAL4* (41). Overexpression of Rala^S25N^ alone did not alter the brain size, while knockdown of *mnb* alone reduced the brain size (Fig. 5K-M). Importantly, combined Rala^S25N^ expression and *mnb* knockdown in the NE resulted in an even smaller brain than *mnb* knockdown alone (Fig. 5N, O). Overall, these results suggest that Mnb and Rala jointly control the development of the wing and the brain, and in the brain their function is required specifically in the NE.

## Discussion

In this work, we have identified a previously unappreciated function of Mnb in the developing *Drosophila* brain. Mnb is required for proper organization of the NE, which harbors precursors of the neural stem cells (NBs), and for maintaining distinct cell fates in the NE and the OPC. By using a strong loss of function allele, *mnb^d419^*, as well as NE-specific knockdowns, we show that loss of *mnb* disrupts NE integrity and compromises the boundary between the NE and the OPC, allowing aberrant expansion of E-cad positive cells into the OPC region. The *mnb* small brain phenotype may therefore derive from increased apoptosis of mis-specified cells, resulting in fewer properly specified NBs and ultimately fewer neurons in the adult brain. Increased levels of apoptosis in *mnb* mutants have been reported previously (21), and we confirmed this observation for the *mnb^d419^* allele (Fig. S3).

We identified a network of in vivo Mnb interacting proteins via AP-MS and uncovered several functional protein groups that may be targeted by or work together with Mnb. Among the proteins in a highly connected endocytosis related cluster, we identified Mnb interactors Reps and Rlip that may form a ternary complex with Mnb. Mnb/DYRK1A and Reps/REPS1/2 physically interact in both *Drosophila* and human cells, suggesting conservation of this interaction and a potential novel regulatory axis for DYRK1A-related diseases such as microcephaly. At the subcellular level, Mnb can recruit Rlip into the cytoplasmic puncta but did not affect Reps localization that was already cytoplasmic (Fig. 3). Beyond physical interactions, Mnb, Reps, and Rlip work together in controlling wing growth, and a ubiquitous joint knockdown of *mnb* and *Rlip* results in lethality, suggesting a common function. Given the functional connection between Rlip and Rala, we investigated a possible interaction between Mnb and Rala, and found that they similarly have a joint function in controlling wing and brain development. In the brain, the functions of Mnb and Rala are required specifically in the NE, suggesting that Rala contributes to Mnb activity in that tissue. Collectively, this evidence establishes a regulatory axis involving Mnb, Reps, Rlip, and Rala (Fig. 6). These proteins likely work together to regulate NE integrity and NB specification during brain development and may be involved in the growth of other tissues such as the wing.

**Figure 6.**
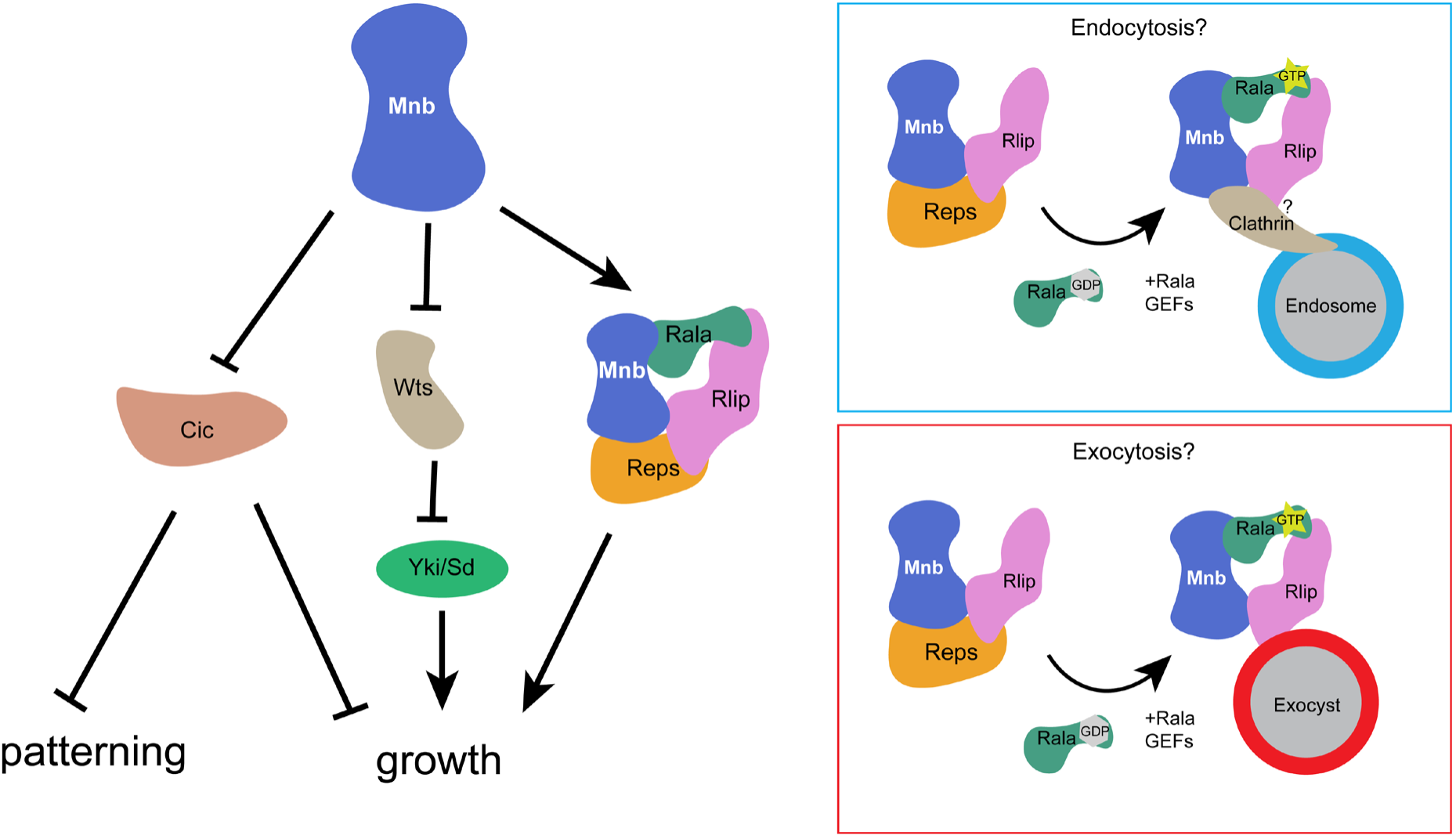
A model for the function of the Mnb/Reps/Rlip/Rala signaling network in development. Left: Mnb participates in at least three regulatory pathways that control organ growth and tissue patterning: ERK/Cic (3), Hippo/Warts (Wts)/Yorkie (Yki) (1), and the Reps/Rlip/Rala pathway described in this work. Right: Mnb may engage the Reps/Rlip/Rala module to control receptor signaling through regulation of endocytosis, exocytosis, or both.

The molecular targets of this regulatory axis are currently unknown. Rala is involved in the regulation of different signaling pathways such as c-Jun N-Terminal Kinase (JNK) (40), EGFR/ERK (42), and Notch (43), in various developmental contexts. The Rala-Rlip-Reps complex has been implicated in both endocytosis and exocytosis of signaling receptors (38, 44). Given the central role for EGFR and Notch signaling in controlling NE development and orchestrating a proper propagation of the proneural wave in the transition to NBs (25, 30, 45, 46), we speculate that the Mnb/Reps/Rlip/Rala signaling network contributes to the regulation of these pathways, possibly at the level of endo-or exocytosis (Fig. 6).

We have shown that Mnb can phosphorylate Reps and that this phosphorylation requires Mnb’s kinase activity, but the significance of this effect is unclear. In S2 cells, the addition of Mnb does not prevent or enhance the Rlip-Reps interaction (Fig. 2). Therefore, we predict that Mnb phosphorylation of Reps has other mechanistic consequences outside of Mnb/Reps/Rlip interactions. It is possible that Mnb-mediated phosphorylation of Reps alters its interactions with Epsins which could affect recruitment of the Reps/Rlip complex to endocytic compartments (47–49), thus regulating signaling. We note that Mnb is likely not the only kinase that phosphorylates Reps, because the Reps bands on a Phos-tag gel are not further slowed down with the addition of Mnb but rather the lower, hypo-phosphorylated forms of Reps gain additional phosphorylations (Fig. 2H). Given the role Reps plays as an adapter protein between Rlip/RALBP1 and Rala in mammalian cells (38), it is possible that it plays a similar role in *Drosophila*. A recent study found that mammalian REPS1 does not interact with RALA but instead helps stabilize RALBP1 until it interacts with RALA, and then Reps dissociates from the RALA-RALBP1 complex (38). The ability of Mnb to phosphorylate Reps and the formation of the Mnb/Reps/Rlip complex may contribute to the dynamics of Reps/Rlip/Rala interactions in flies. Such dynamics may be critical for the proper regulation of signaling pathways responsible for the NE to NB transition.

We have previously established that Mnb contributes to the control of organ growth and tissue patterning in *Drosophila* via downregulation of the Hippo pathway kinase Warts (1) and via inactivation of the transcriptional repressor Capicua in parallel with ERK (3). Our current study delineates a third important signaling function of Mnb that involves trafficking regulators Reps, Rlip, and Rala (Fig. 6). In the future, it would be of interest to determine how these pathways are coordinated to mediate proper development of the brain and other tissues.

## Materials and Methods

### Fly stocks

All *Drosophila* stocks were maintained on standard yeast-cornmeal-agar medium at 25 °C or 18 °C as indicated. The following fly lines were used in this study: *mnb^d419^* (5), *UAS-wRNAi* (Bloomington #33762), *UAS-mnbRNAi* (Kyoto #7826R-3), *da-GAL4* (50), *c855a-GAL4* (Bloomington #6990) (41), *MS1096-GAL4* (51), *UAS-RepsRNAi1* (VDRC # 24719), *UAS-RepsRNAi2* (VDRC # 110704), *UAS-RlipRNAi* (VDRC #105976), *UAS-Rala*^S25N^ (Bloomington # 32094), *Dp(1;3)DC337* (Bloomington # 30441), *Dp(1;3)DC033* (Bloomington # 38493).

### Immunostaining and fluorescence microscopy

3^rd^ instar larval brains were dissected in 1x PBS with 0.3% Triton X-100 (Sigma Aldrich cat#T9284) and fixed in 4% paraformaldehyde (Electron Microscopy Sciences cat#15710) in 1x PBS for 20 minutes at room temperature (RT) in darkness. Brains were washed three times for 5 minutes at RT with 1x PBS before blocking for 30 minutes with 5% normal goat serum (Fisher cat#50-413-405) in 1x PBS with Triton X-100. Brains were incubated in primary antibody (rat anti D-Cad2, DSHB, 1:20, rabbit anti cleaved Dcp-1, CST cat#9578, 1:100, or rat anti Miranda, Abcam CD#5-7E9BG5AF4, 1:500) diluted in 1x PBS 0.3% Triton X-100 overnight. To immunostain two genotypes in the same well, eye discs and the mouth hooks were removed from brains from one genotype and were kept attached in the other genotype. Following primary antibody incubation, brains were washed 3 times for 20 minutes with 1X PBS before incubation with secondary antibody (goat anti rat Alexa Fluor 555, Thermo cat#A-21434, 1:500 or goat anti rabbit Alexa Flour 555, Thermo cat#PIA32732) diluted in 1X PBS 0.3% Triton X-100 for 3 hours. Brains were washed following secondary antibody 3 times for 20 minutes with 1X PBS before 10-15 brains per slide were mounted between two coverslips placed approximately 0.5 cm apart in 15 µL of ProLong Gold Antifade Mountant, With DAPI (Fisher cat#P36931).

Fluorescent images were acquired using a Zeiss LSM 880 confocal microscope at 40x objective with 1 Airy Unit pinhole. Z-stacks were acquired with optimal slice numbers and maximum-intensity projections were performed in Zen Blue software. Arrowheads and neuroepithelium-neuroblast boundary lines were added in Adobe Illustrator 2023. Corrected total fluorescence was calculated as: raw intensity of Dcp-1 in optic lobe – (average background intensity × optic lobe area) in Fiji (52).

### Affinity purification from embryos and mass spectrometry analysis

Flies from the Mnb-tRFP line and *yw* controls were set up in 5 L fly cages with apple juice plates. Flies were allowed to lay eggs for 15 hours at RT, then apple juice plates containing embryos were aged for 3 hours at 25 °C. Approximately 1 g of embryos were dechorionated with 50% bleach for 1.5 minutes and washed with water. Embryos were then lysed on ice with 4 mL of ice-cold default lysis buffer (DLB) (50 mM Tris (pH 7.5), 5% glycerol, 0.2% IGEPAL, 1.5 mM MgCl_2_, 125 mM NaCl, 25 mM NaF, 1 mM Na_3_VO_4_, with 2x cOmplete Protease Inhibitor, MilliporeSigma cat#11697498001, 1 tablet per 25 mL of lysis buffer) with additional IGEPAL added to a final concentration of 0.5%, in a glass homogenizer using 30 strokes with a tight pestle. The lysates were incubated on ice for 20 minutes and then centrifuged at 16,000 rcf for 20 minutes at 4 °C. The supernatant was then filtered through a pre-chilled 0.45 µm filter (Fisher cat#0975421). The filtered supernatant was then incubated with 50 µL of packed Pierce Control Agarose beads (Thermo cat#26150) at 4 °C for 30 minutes with rotation. The lysate was then incubated with 20 µL of packed RFP-trap Agarose beads (Bulldog Bio cat#RTA020) for 2 hours at 4 °C with rotation. After binding, the beads were washed 5 times with 1 mL of DLB with 0.5% IGEPAL. Then 40 µL of 4x SDS sample buffer were added to beads and the samples were heated at 95 °C for 6 minutes. Samples were analyzed on a NuPAGE 4-12% Bis-Tris Protein Gel (Fisher cat#NP0335) and SilverQuest Silver Staining Kit (Fisher cat#LC6070) to confirm sample quality. Protein samples for mass spectrometry were prepared by running 1 cm into an 8% Tris-Glycine SDS-PAGE gel followed by Coomassie blue staining. Two gel pieces (>75 kDa and <75 kDa) were cut from the gel for each sample and sent for mass spectrometry (MS) analysis. The MS analysis was conducted on two biological replicates for experimental samples (Mnb-tRFP) and three control samples (*yw*).

Samples were analyzed by liquid chromatography/tandem mass spectrometry (nanoLC-MS/MS) using Thermo Scientific Orbitrap mass spectrometer by the Taplin Mass Spectrometry Facility at Harvard Medical School. Mass spectrometry data were analyzed using Sequest and searches were run using a database of annotated *Drosophila* proteins, with the following search parameters: mass tolerance = 2, mass units = amu, fragment ion tolerance = 1, ion series = nB;nY;b;y, Mods = 15.9949146221 M 14.0157 C, enzyme = trypsin. MS results from two gel pieces for each sample were combined by summing unique peptide counts for each protein. The results for experimental and control samples were compared using the Significance Analysis of INTeractome (SAINT)(v2.5.0) using peptide counts identified for each protein. Any protein with the SAINT score (AvgP) above 0.8 was considered a high-confidence interactor and was included in the interactome. A complete SAINT output is provided in Dataset S1.

The Mnb protein interactome was generated using the STRING database app in Cytoscape using the following standard options: Network type: Full STRING Network, Confidence Score (Cutoff): 0.40, Maximum additional interactors: 0, Use smart delimiters: checked. The resulting STRING network was clustered using the clusterMaker MCL Cluster app in Cytoscape. The following MCL cluster settings were used: Granularity parameter: 2.5, Array Source: None, Edge cut off: 0, Assume edges are undirected: check, Adjust loops before clustering: check, Create new clustered network: check. To analyze Gene Ontology of the resulting clustered network, highly interconnected clusters were analyzed using STRING enrichment.

### Plasmid construction

pMT-Reps-Flag and pMT-Reps-HA were generated by cloning Reps-RA isoform from FMO04419 (DGRC Stock 1612020) into pMT-V5-His vector (Invitrogen cat#V412020). pMT-Mnb-V5 was generated by cloning Mnb-RH isoform from FMO12028 (DGRC Stock 1639183) into pMT-V5-His vector. pMT-Rlip-HA was generated by cloning the Rlip-RA isoform from GH01995 (DGRC Stock 12147) into pMT-V5-His vector. pMT-mCherry-Rlip and pMT-Rlip-mCherry were cloned from pMT-Rlip-HA using NEB HiFi Assembly kit (NEB cat#E5520) to exclude the retained intron and incorporate the mCherry open reading frame into pMT-V5-His vector. pcDNA3.1-DYRK1A-V5 was generated by cloning DYRK1A from pMH-SFB-DYRK1A (Addgene cat#101770) into pcDNA3.1(-) (ThermoFisher cat#V79520). pcDNA3.1-REPS2-Flag was obtained from Sinobiological (cat#HG20820-NF). pcDNA3.1-REPS1-Flag was generated by cloning REPS1 from pcMV-REPS1 (Sinobiological cat#HG22238-UT) into pcDNA3.1(-).

### Co-immunoprecipitation (co-IP)

*Drosophila* S2 cells were transfected with the indicated plasmids or blank pMT-V5-His vector (Invitrogen cat#V412020) using Effectene transfection reagent (Qiagen cat#301427). 24 hours after transfection, cells were induced with 0.35 mM CuSO_4_ overnight at 25 °C. Cells were then collected, washed with 10 mL of 1x PBS (Fisher cat#BP399), and lysed on ice for 20 minutes with 700 µL of ice-cold DLB. The lysate was centrifuged at 20,000 rcf for 20 mins at 4 °C. To generate lysate samples, 50 µL of lysate were added to 25 µL of 4X SDS sample buffer and heated for 5 mins at 95 °C. 450 µL of the lysate were incubated with 20 µL of packed anti-V5 beads (MilliporeSigma cat#A7345), anti-Flag beads (MilliporeSigma cat#F2426), anti-HA beads (MilliporeSigma cat#E6779), or anti-RFP beads (Bulldog Bio cat#RTA020) for 2 hours at 4°C. The beads were then washed 5 times with 1 mL of DLB. 40 µL of 4X SDS sample buffer was added and the beads were incubated at 95 °C for 5 minutes to generate IP samples. The lysate and IP samples were run on SDS-PAGE followed by western blot.

### Phos-tag analysis and western blotting

Samples were prepared using EDTA-free DLB (composition as above, except with 2x cOmplete EDTA-free Protease Inhibitor, MilliporeSigma cat#11873580001, 1 tablet per 25 mL of lysis buffer) and immunoprecipitated with anti-Flag beads. IP sample was analyzed by standard SDS-PAGE and Phos-tag analysis. A 6% Phos-tag gel with 50 µM Phos-tag (FujiFilm Wako Chemicals cat#304-93526) and 100 µM MnCl_2_ (Fisher cat#205891000) was run at 80 V for 4 hours at 4 °C. Following electrophoresis, the Phos-tag gel was washed with 10 mM EDTA in 1x Transfer Buffer 2 times for 10 minutes followed by one wash with 1x Transfer Buffer for 10 minutes. The gel was then wet transferred at 90 V for 2 hours at 4 °C. The blots were then blocked in blocking buffer (LI-COR cat#927-70001) for 30 minutes before incubation with primary antibody (rabbit anti-FLAG, Sigma-Aldrich cat#F7425, 1:1000) and secondary antibody donkey anti-rabbit IgG (1:20,000, LI-COR cat#926-32213). Blots were scanned on LI-COR Odyssey CLx Imaging System. Other antibodies used for western blotting were mouse anti-V5 (MilliporeSigma cat#V8012, 1:1000), rabbit anti-HA (MilliporeSigma cat#H6908, 1:1000), rabbit anti-dsRed (Takara cat#632496, 1:1000), and Goat anti-Mouse IgG (LI-COR cat#926-68070, 1:20,000).

### Wing and brain analysis

Adult female wings were removed and mounted in 3:1 CMCP-10 Mounting Medium (Fisher cat#50-190-0430)/lactic acid (Fisher cat#A162). Wing images were taken with a Zeiss Axiocam 712 camera on an Olympus BX60 Microscope. The wing size was calculated as the wing area and analyzed using Fiji. Adult female heads were removed and fixed in 4% paraformaldehyde in 1x PBS for 20 mins at RT in darkness. Following fixation, heads were washed 3 times in 1x PBS for 5 minutes at RT. Brains were dissected from fixed heads before mounting. 10-15 brains per slide were mounted between two coverslips placed approximately 0.5 cm apart in 15 µL of ProLong Gold Antifade Mountant, With DAPI (Fisher cat#P36931). Slides were stored overnight in darkness at RT before sealing the edges with clear nail polish. Fluorescent images of brains were taken with a Zeiss Axiocam 712 camera on an Olympus BX60 Microscope with a 10x objective. Brain size was measured in Fiji as the area of the DAPI positive brain tissue.

## Acknowledgements

We thank the Bloomington *Drosophila* Stock Center, *Drosophila* Genomics Resource Center, Vienna *Drosophila* Resource Centre, Kyoto Drosophila Stock Center, and Developmental Studies Hybridoma Bank for their services; and Alexandra Sheydvasser for help with the cloning of REPS1/2 constructs. Mass spectrometry was performed at the Taplin Mass Spectrometry Facility at Harvard Medical School. This work was supported by the NIH grant GM141843 to A.V. M.B. was supported by the University of Massachusetts Boston Doctoral Fellowship, E.S. was supported by the University of Massachusetts Boston College of Science and Mathematics Undergraduate Research Fellowship. We thank Nathan Strozewski for critical reading of the manuscript.

---

**Supplemental Figure S1.**
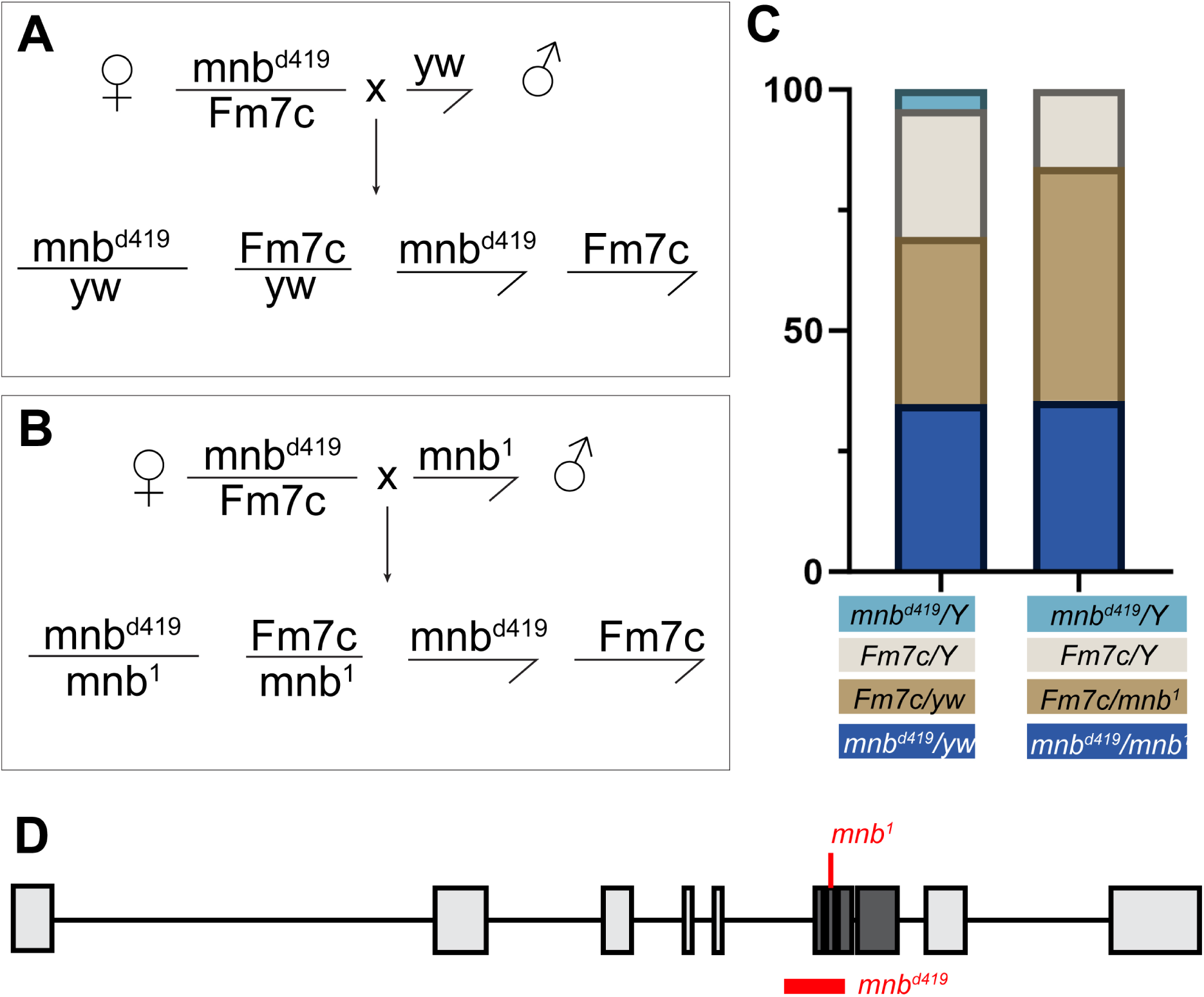
*mnb^1^* can complement *mnb^d419^*: (A) Crossing scheme and expected progeny of *mnb^d419^* females crossed with *yw* males. (B) Crossing scheme and expected progeny of *mnb^d419^* females crossed with *mnb^1^* males. (C) Distribution of observed progeny from crosses in A-B. (D) The nature of the *mnb^d419^* allele (modified from (5)). Mnb exons are in light grey, kinase domain in dark grey. The *mnb^1^* point mutation and the imprecise deletion in *mnb^d419^* that deletes half of the kinase domain are marked in red.

**Supplemental Figure S2.**
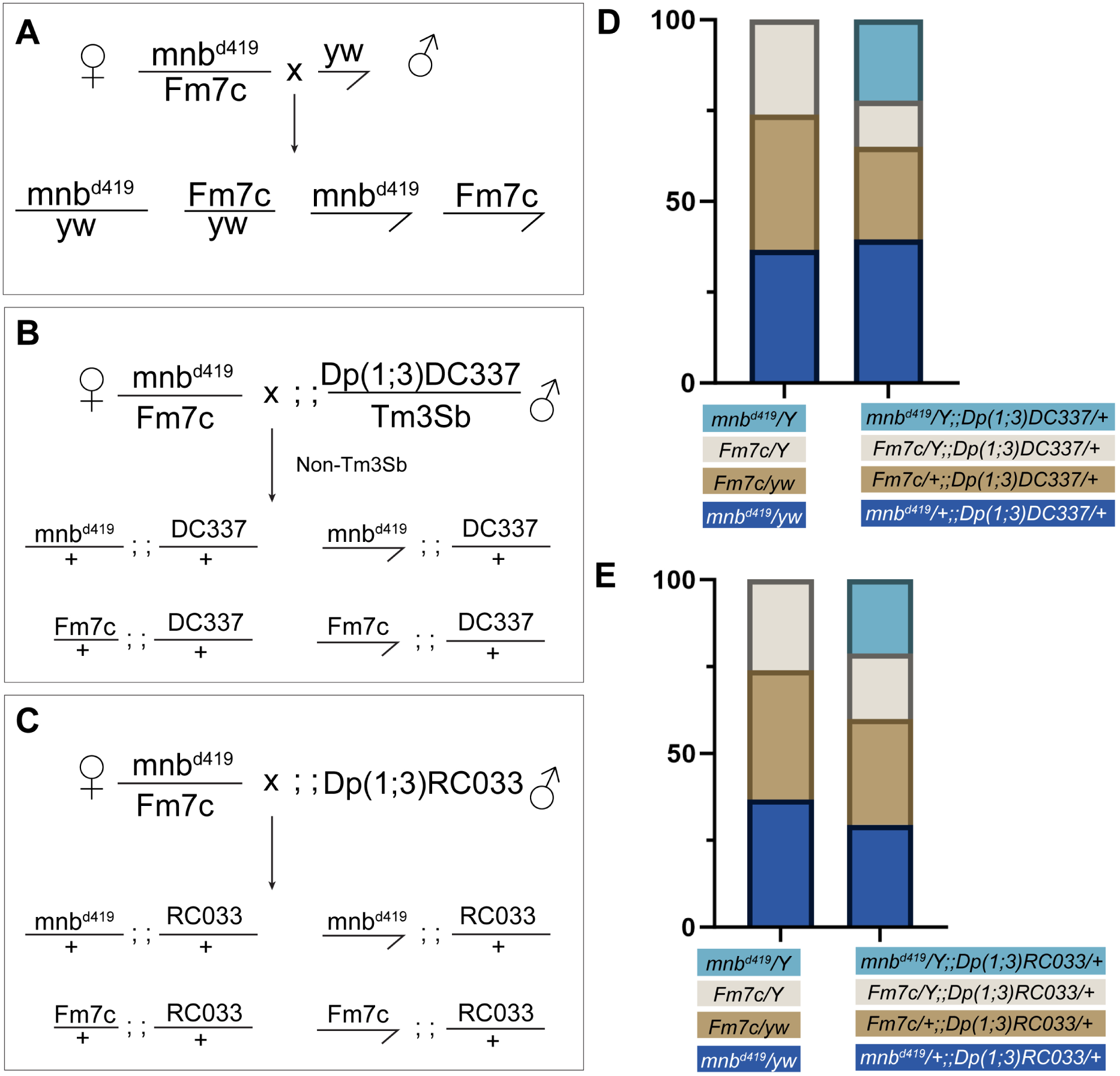
Duplication lines can complement *mnb^d419^*: (A) Crossing scheme and expected progeny of *mnb^d419^* females crossed with *yw* males. (B) Crossing scheme and expected progeny of *mnb^d419^* females crossed with *Dp(1;3)DC337-30441* males. (C) Crossing scheme and expected progeny of *mnb^d419^* females crossed with *Dp(1;3)DC033-38493* males. (D) Distribution of observed progeny from crosses in (A-B). (E) Distribution of observed progeny from crosses in (A, C). Both duplications rescued the lethality associated with *mnb^d419^*.

**Supplemental Figure S3.**
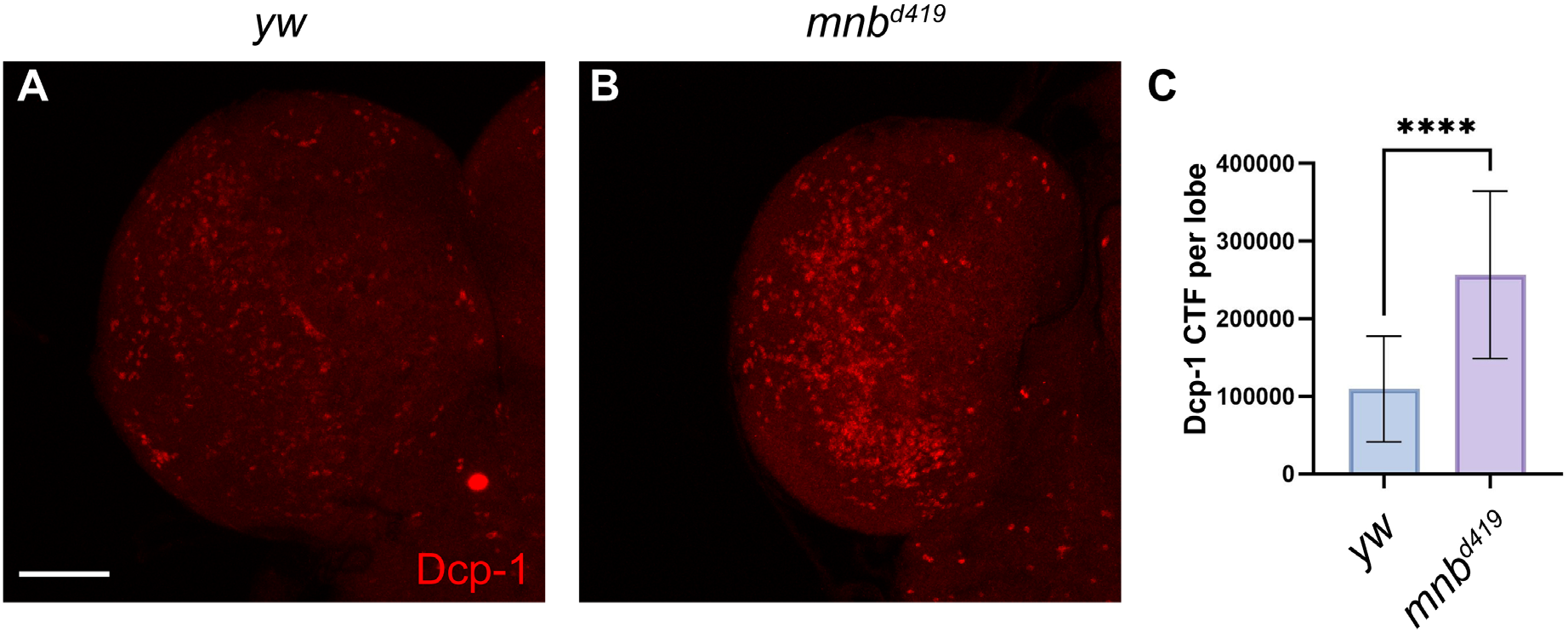
Mnb is required for limiting apoptosis. (A-B) Confocal maximum intensity projections of brains from control *yw* (A) and *mnb^d419^* (B) 3^rd^ instar larvae immunostained for cleaved death caspase 1, Dcp-1 (shown in red). Scale bar, 50 µm. (C) Quantification of cleaved Dcp-1 corrected total fluorescence per optic lobe shown in (A-B). n=14, \*\*\*\**p*<0.0001, *p* value calculated using Student’s t-test. Error bars indicate standard deviation.

**Supplemental Figure S4.**
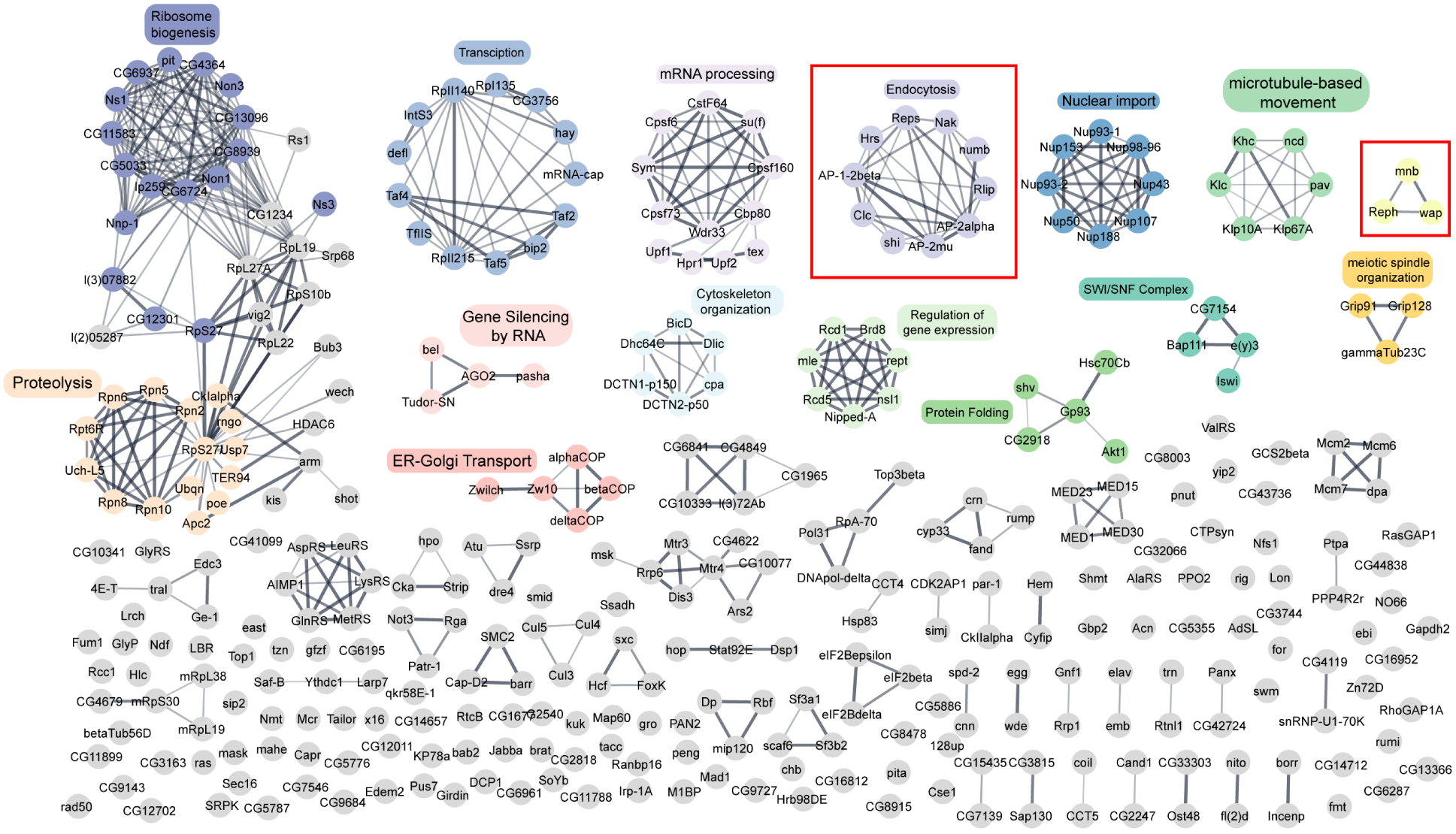
Mnb protein interaction network: Significantly enriched proteins identified in affinity purification-mass spectrometry (AP-MS) of *Drosophila* embryos expressing endogenously tagged Mnb. All proteins in this network were identified as high-confidence Mnb interactors in this analysis. Edges in this network were derived from the STRING protein database, and clusters were generated using the clusterMaker MCL Cluster app in Cytoscape. Gene Ontology analysis revealed biological process or cellular component enrichment terms, indicated near the respective clusters. Ribosome biogenesis, FDR=8.01E-21; proteolysis, FDR=9.25E-9; transcription, FDR=5.7E-12; mRNA processing, FDR=1.78E-8; endocytosis, FDR=1.64E-8; nuclear import, FDR=4.9E-13; microtubule-based movement FDR=6.6E-8; meiotic spindle organization, FDR=3.6E-5; SWI/SNF complex, FDR 2.66E-7; protein folding, FDR=3.9E-4; regulation of gene expression, FDR=4.4E-4; cytoskeleton organization, FDR=5.57E-6; ER-golgi transport, FDR=2.3E-5; gene silencing, FDR=3.04E-6.

**Supplemental Figure S5.**
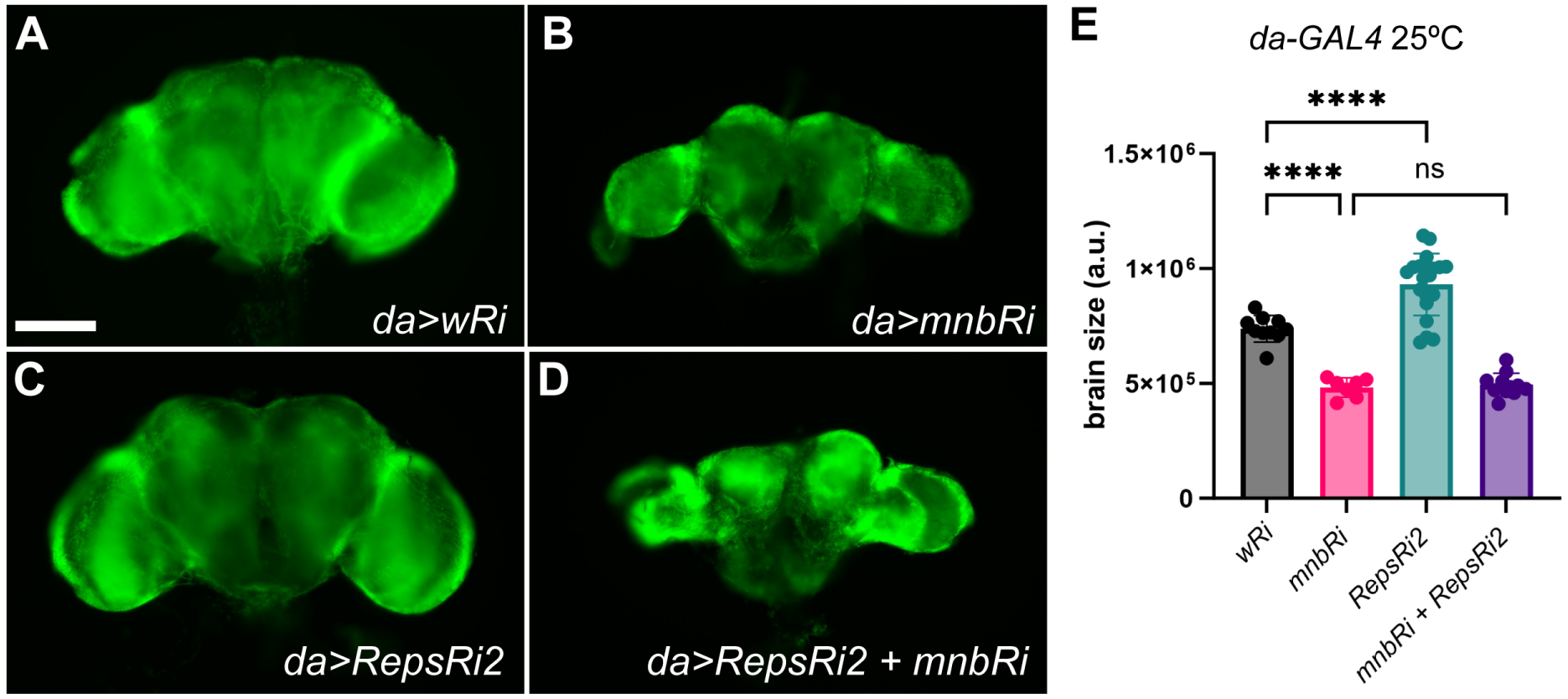
(A-D) Adult brains from female flies expressing the indicated RNAi transgenes using *da-GAL4*. (E) Quantification of brain size shown in (A-D) (n≥7 for each genotype). Brain size was measured as total area. DAPI signal shown in green. \*\*\*\**p*<0.0001, *p* value calculated using ANOVA. Error bars represent SD. Scale bar, 100 µm.

